# SMURF2 phosphorylation at Thr249 modifies the stemness and tumorigenicity of glioma stem cells by regulating TGF-β receptor stability

**DOI:** 10.1101/2021.04.27.441592

**Authors:** Manami Hiraiwa, Kazuya Fukasawa, Takashi Iezaki, Hemragul Sabit, Tetsuhiro Horie, Kazuya Tokumura, Sayuki Iwahashi, Misato Murata, Masaki Kobayashi, Gyujin Park, Katsuyuki Kaneda, Tomoki Todo, Atsushi Hirao, Mitsutoshi Nakada, Eiichi Hinoi

## Abstract

Glioma stem cells (GSCs) contribute to the pathogenesis of glioblastoma, the most malignant form of glioma. The implication and underlying mechanisms of SMAD specific E3 ubiquitin protein ligase 2 (SMURF2) on the GSC phenotypes remain unknown. We previously demonstrated that SMURF2 phosphorylation at Thr^249^ (SMURF2^Thr249^) activates its E3 ubiquitin ligase activity. Here, we demonstrate that SMURF2^Thr249^ phosphorylation plays an essential role in maintaining GSC stemness and tumorigenicity. *SMURF2* silencing augmented the self-renewal potential and tumorgenicity of patient-derived GSCs. The SMURF2^Thr249^ phosphorylation level was low in human glioblastoma pathology specimens. Introduction of the *SMURF2*^T249A^ mutant resulted in increased stemness and tumorgenicity of GSCs, recapitulating the *SMURF2* silencing. Moreover, the inactivation of SMURF2^Thr249^ phosphorylation increases TGF-β receptor (TGFBR) protein stability. Indeed, *TGFBR1* knockdown markedly counteracted the GSC phenotypes by *SMURF2*^T249A^ mutant. These findings highlight the importance of SMURF2^Thr249^ phosphorylation in maintaining GSC phenotypes, thereby demonstrating a potential target for GSC-directed therapy.

## Introduction

SMAD specific E3 ubiquitin protein ligase 2 (SMURF2) is the E3 ubiquitin ligase responsible for specifying the substrates for ubiquitination and degradation by proteasomes (1, 2). Accumulating evidence indicates SMURF2 regulates a wide array of physiological processes, including cell proliferation, invasion, self-renewal, and migration, through its regulation of a variety of signaling pathways (3–5). The E3 ubiquitin ligase activity of SMURF2 is regulated at the post-transcriptional level through SUMOylation, methylation, and phosphorylation (6–8), as well as at the transcriptional level (9). We recently demonstrated that the phosphorylation of SMURF2 at Thr^249^ (SMURF2^Thr249^) by extracellular signal-regulated kinase 5 (ERK5) plays an essential role in maintaining the stemness of mesenchymal stem cells (MSCs), which contributes to skeletogenesis (10). Mechanistically, SMURF2^Thr249^ phosphorylation activates its E3 ubiquitin ligase activity, which modifies the stability of SMAD proteins, which in turn transcriptionally activate the expression of SOX9, the principal transcription factor of skeletogenesis in MSCs.

Gliomas, which represent approximately 80% of all primary malignant brain tumors in humans, can be categorized into four grades according to the World Health Organization (WHO) classification criteria: grade I, grade II, grade III, and grade IV (glioblastoma, GBM) (11, 12). GBM, the most malignant form of glioma, is one of the most aggressive and deadly types of cancer. Patients with GBM have a very poor prognosis, with a five-year survival rate of only 5.1% (13, 14). Glioma stem cells (GSCs), also known as glioma-initiating cells, are a subpopulation of tumor cells that exhibit stem cell-like capacities such as self-renewal and tumor-initiating capacities (15–17). Recent studies have determined that GSCs contribute to high rates of therapeutic resistance and rapid recurrence (18, 19), cancer invasion, immune evasion, tumor angiogenesis, and the recruitment of tumor-associated macrophages, which indicates that targeting GSCs is an efficacious strategy for improving GBM treatment (20–22).

Transforming growth factor-β (TGF-β) signaling, which is tightly regulated through protein ubiquitination (23, 24), has been shown to play a crucial role in maintaining the stemness and tumorigenicity of GSCs through several pathways including the SMAD-SOX4-SOX2 axis and the SMAD-LIF-JAK-STAT pathway (25, 26). SMAD7 acts as a scaffold protein to recruit SMURF2 to the TGF-β receptor (TGFBR) complex to facilitate its ubiquitination (27). This leads to the proteasome-mediated degradation of TGFBRs and the attenuation of TGF-β signaling. Ubiquitin-specific peptidase 15 (USP15), a deubiquitinating enzyme, binds to the SMAD7-SMURF2 complex and deubiquitinates and stabilizes TGFBR1, resulting in enhanced TGF-β signaling (28, 29). The balance between USP15 and SMURF2 activities determines the activity of TGF-β signaling and subsequent oncogenesis in GBM. Indeed, a deficiency in USP15 decreases the oncogenic capacity of GSCs due to the repression of TGF-β signaling (28); conversely, USP15 amplification confers poor prognosis in individuals with GBM (30). However, although SMURF2 should be assumed to play an opposite role from that of USP15, no reports have yet directly addressed the implication and underlying mechanisms of SMURF2 on the GSC phenotypes and subsequent glioma pathogenesis both *in vivo* and *in vitro*.

In this study, we reveal that *SMURF2* silencing by shRNA resulted in an augmentation of the self-renewal potential and tumorigenicity of GSCs. The SMURF2^Thr249^ phosphorylation level was downregulated in GBM patients, regardless of the lack of marked changes in its mRNA and protein levels. Additionally, the SMURF2^Thr249^ phosphorylation level was lower in GSCs than that in differentiated glioma cells. The inactivation of SMURF2^Thr249^ phosphorylation by a non-phosphorylatable mutant (*SMURF2*^T249A^ mutant) increased the self-renewal potential and tumorigenicity of GSCs, thus mimicking the GSC phenotype in *SMURF2* silencing. Mechanistically, SMURF2^Thr249^ phosphorylation activates its E3 ubiquitin ligase activity, which decreases the protein stability of TGFBR1 via proteasome-mediated degradation. Finally, *TGFBR1* silencing rescues the increased self-renewal potential and tumorigenicity of GSCs by inactivating SMURF2^Thr249^ phosphorylation. Collectively, these findings highlight the importance of SMURF2^Thr249^ phosphorylation in maintaining the stemness and tumorigenicity of GSCs; these findings also indicate that SMURF2^Thr249^ phosphorylation could be an important posttranslational modification in treatment strategies aimed at disrupting GSCs.

## Results

### Targeting *SMURF2* promotes the self-renewal potential and tumorigenicity of GSCs

We first elucidated the functional significance of SMURF2 in maintaining GSCs *in vitro* by targeting *SMURF2* expression using lentiviral shRNA (sh*SMURF2*) in TGS-01 and TGS-04 GSCs, which are human GBM patient-derived GSCs. Disruption of *SMURF2* with shRNA significantly increased GSC tumorsphere formation in both TGS-01 and TGS-04 GSCs (Fig. 1A). Additionally, an *in vitro* limiting dilution assay demonstrated that the self-renewal potential of GSCs was significantly increased by *SMURF2* silencing in both TGS-01 and TGS-04 GSCs (Fig. 1B). Furthermore, *SMURF2* knockdown resulted in the significant upregulation of the stem cell transcription factors SOX2 and SOX4 in TGS-01 GSCs, along with a marked reduction in the SMURF2 protein level (Fig. 1C). Conversely, disrupting *SMURF2* did not significantly alter cell apoptosis in TGS-01 GSCs (Fig. 1D).

**Figure 1.**
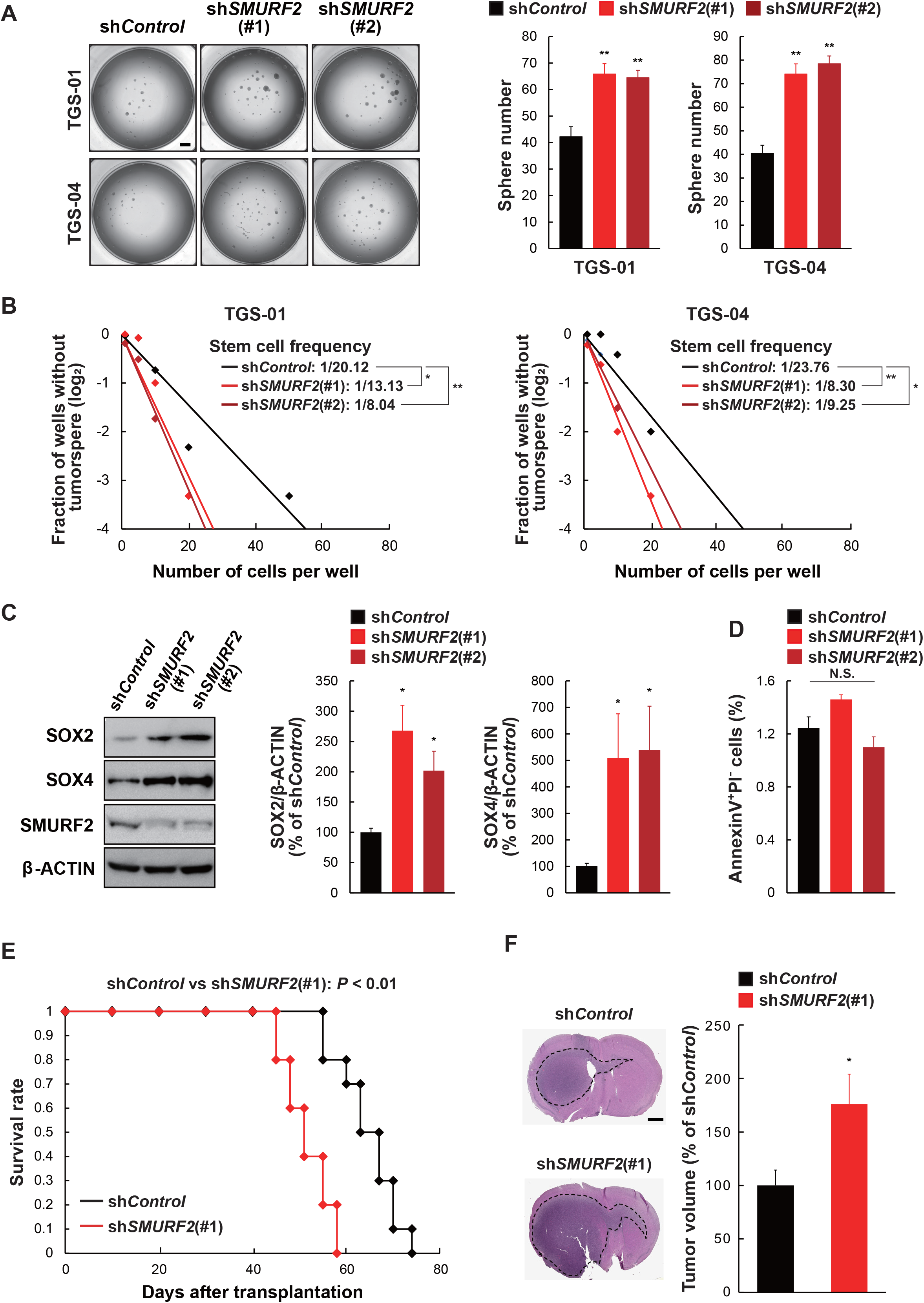
*SMURF2* silencing promotes tumor growth and self-renewal of GSCs. TGS-01 and TGS-04 GSCs were infected with sh*SMURF2* (#1 and #2), followed by determination of (A) tumorsphere number (*n*=8), (B) stem cell frequency by *in vitro* limiting dilution assay (estimated frequencies of clonogenic cells in GSC tumorsphere were calculated by ELDA analysis), (C) protein levels of SOX2, SOX4, and SMURF2; β-ACTIN served as a loading control (*n*=3), and (D) cell apoptosis (*n*=3). (E) Development of gliomas after intracranial transplantation of sh*SMURF2*-infected TGS-01 GSCs. Survival of mice was evaluated by Kaplan-Meier analysis (*n*=10). *P* value was calculated using a log-rank test. (F) Histological analyses of brains dissected at 30 days after intracranial transplantation. Tissue sections were stained with H&E (*n*=5). \**P* < 0.05, \*\**P* < 0.01, significantly different from the value obtained in cells infected with sh*Control*. N.S., not significant. Values are expressed as the mean ± S.E. and statistical significance was determined using (A and C) the one-way ANOVA using the Bonferroni *post hoc* test, and (F) Student’s *t*-test. Scale bar: 1 mm.

We next examined whether *SMURF2* silencing could affect the tumorigenic potential of GSCs in an orthotopic xenograft mouse model. Equal numbers of TGS-01 GSCs transduced with either sh*SMURF2* or sh*Control* were intracranially injected into immunocompromised mice. The mice inoculated with the sh*SMURF2*-infected TGS-01 GSCs had a significantly shortened survival compared with the mice injected with the sh*Control*-infected cells (Fig. 1E). Moreover, the histological examination demonstrated that the mice inoculated with sh*SMURF2*-infected TGS-01 GSCs displayed larger tumors compared with the mice injected with sh*Control*-infected cells (Fig. 1F). Collectively, our findings in patient-derived GSCs *in vitro* and in the *in vivo* orthotopic xenograft model indicate the importance of SMURF2 in the self-renewal potential and tumorigenicity of GSCs.

### SMURF2^Thr249^ phosphorylation level is lower in human GBM tissues and human GBM patient-derived GSCs

We next assessed whether our findings were relevant to clinical data in glioma patients using publicly available datasets and our clinical samples. No marked alterations of *SMURF2* mRNA levels were found among grades II, III, and IV cancer or among classical, mesenchymal, and proneural tumors, according to the Cancer Genome Atlas (TCGA) (Fig. 2A). Moreover, in accordance with the lack of marked alterations of the *SMURF2* mRNA levels in glioma specimens in the TCGA database, we confirmed that the SMURF2 protein level was comparable between control nonneoplastic brain tissue (NB), diffuse astrocytoma (grade II), anaplastic astrocytoma (grade III), and GBM (grade IV) in our clinical samples (Fig. 2B and 2C).

**Figure 2.**
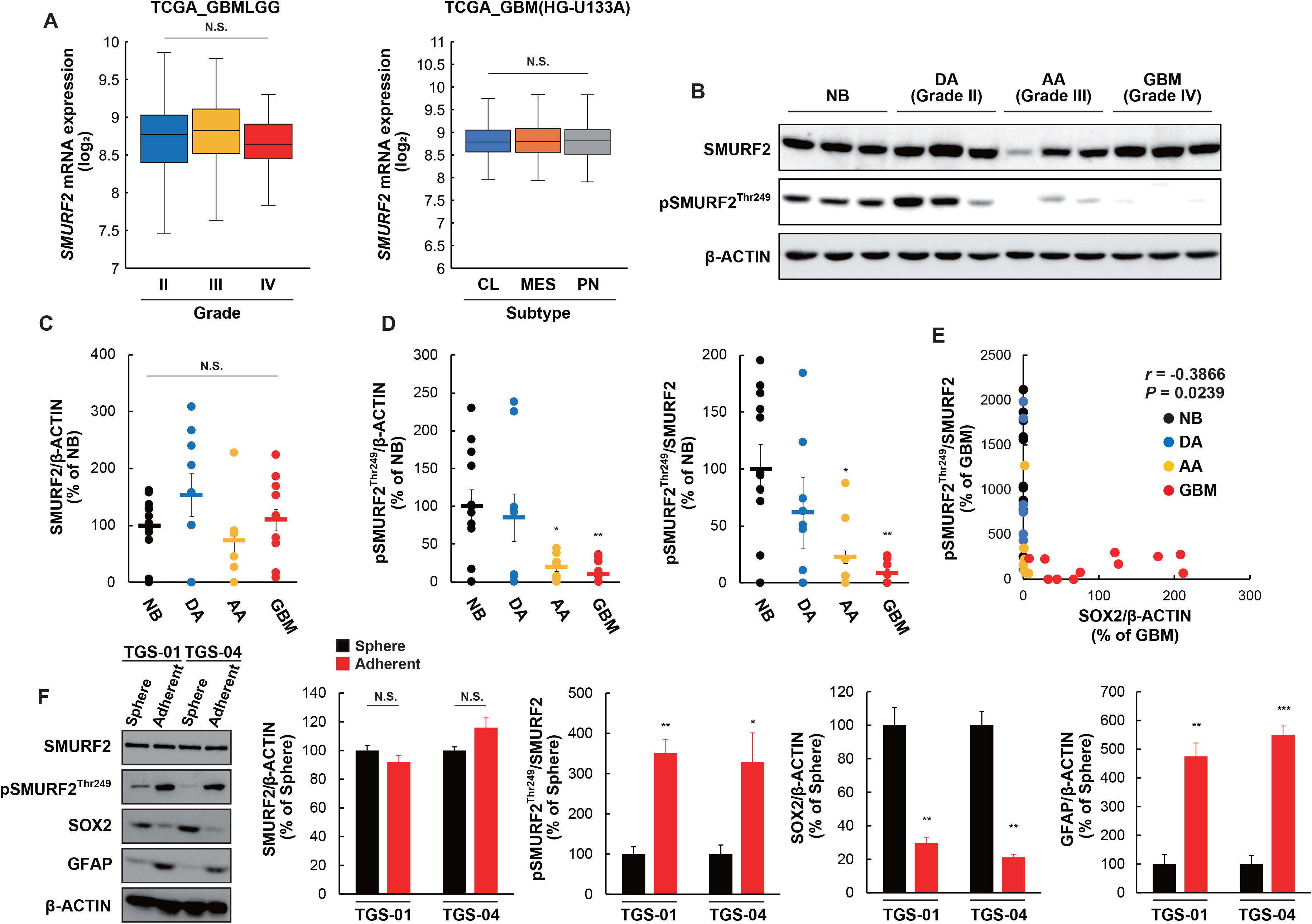
SMURF2^Thr249^ phosphorylation is decreased in anaplastic astrocytoma and GBM specimens, and is a negative correlation with stem cell marker. (A) mRNA expression of *SMURF2* in each grade (grade II, *n*=226; grade III, *n*=244; grade IV, *n*=150) or subtype (classical (CL), *n*=199; mesenchymal (MES), *n*=166; proneural (PN), *n*=163) of glioma. The data was obtained and analyzed using GlioVis database. (B-D) Determination of protein levels of SMURF2 and pSMURF2^Thr249^ in human glioma samples. Nonneoplastic brain tissue (NB) (*n*=12), diffuse astrocytoma (DA) Grade II (*n*=9), anaplastic astrocytoma (AA) Grade III (*n*=9), glioblastoma (GBM) Grade IV (*n*=16). (E) Correlation between SOX2 and pSMURF2^Thr249^ in glioma samples. \**P* < 0.05, \*\**P* < 0.01, significantly different from the value obtained in NB. (F) TGS-01 and TGS-04 cells were cultured in neurosphere medium or adherent culture medium, followed by determination of protein levels of SOX2, GFAP, SMURF2 and pSMURF2^Thr249^; β-ACTIN served as a loading control (*n*=3). \**P* < 0.05, \*\**P* < 0.01, \*\*\**P* < 0.001, significantly different from the value obtained in Sphere. N.S., not significant. Values are expressed as the mean ± S.E. and statistical significance was determined using (A) Tukey’s Honest Significant Difference test, (C and D) the one-way ANOVA using the Bonferroni *post hoc* test, and (F) Student’s *t*-test. *r*, Pearson’s correlation coefficient.

These results led us to investigate whether the posttranslational modification of SMURF2 could be modified in human glioma specimens to reveal the functional importance of SMURF2 in the development and progression of gliomas. Given that our previous study reported that SMURF2^Thr249^ phosphorylation plays an essential role in maintaining the stemness of MSCs (10), we next examined the SMURF2^Thr249^ phosphorylation level in human glioma specimens. The SMURF2^Thr249^ phosphorylation level was significantly lower in the GBM (grade IV) and anaplastic astrocytoma (grade III) specimens than in NB specimens (Fig. 2B and 2D). Moreover, the SMURF2^Thr249^ phosphorylation level was negatively correlated with the protein level of SOX2, a stem cell transcription factor (31, 32), in glioma specimens (Fig. 2E).

Further, we compared the SMURF2^Thr249^ phosphorylation level in GSCs and that in differentiated glioma cells. For this, TGS-01 and TGS-04 cells were cultured in neurosphere culture condition (for GSCs) or adherent culture condition (for differentiated glioma cells). Under neurosphere culture condition, TGS-01 and TGS-04 GSCs displayed a significant lower SMURF2^Thr249^ phosphorylation level, in addition to a higher SOX2 level and a lower GFAP level, when compared with TGS-01 and TGS-04 cells cultured under adherent culture condition (Fig. 2F). Conversely, SMURF2 protein level was comparable between cells under the two culture conditions (Fig. 2F). Therefore, these results indicated that the SMURF2^Thr249^ phosphorylation level was significantly lower in GSCs than that in differentiated glioma cells. Our experimental findings aligned with publicly available clinical data suggest that SMURF2^Thr249^ phosphorylation rather than SMURF2 levels (protein and mRNA) might be associated with tumor grade and glioma stemness in humans. Thus, SMURF2^Thr249^ phosphorylation may serve as a prognostic marker of GBM.

### SMURF2^Thr249^ phosphorylation is implicated in the self-renewal potential and tumorigenicity of GSCs

We next determined whether the SMURF2^Thr249^ phosphorylation is implicated in the maintenance of GSCs *in vitro*. To this end, a T249A *SMURF2* mutant construct (hereafter referred to as *SMURF2*^T249A^), in which threonine was replaced by alanine to prevent phosphorylation, was lentivirally infected in both TGS-01 and TGS-04 GSCs. The introduction of *SMURF2*^T249A^ significantly increased tumorsphere formation and the self-renewal potential in both TGS-01 and TGS-04 GSCs; conversely, these changes were significantly decreased after the introduction of wild-type SMURF2 (hereafter referred to as *SMURF2*^WT^) (Fig. 3A and 3B). Additionally, an immunoblotting analysis revealed that the protein levels of SOX2 and SOX4 were significantly upregulated by *SMURF2*^T249A^ but significantly downregulated by *SMURF2*^WT^ (Fig. 3C). Conversely, cell apoptosis was not markedly altered by either *SMURF2*^T249A^ or *SMURF2*^WT^ in TGS-01 GSCs (Fig. 3D).

**Figure 3.**
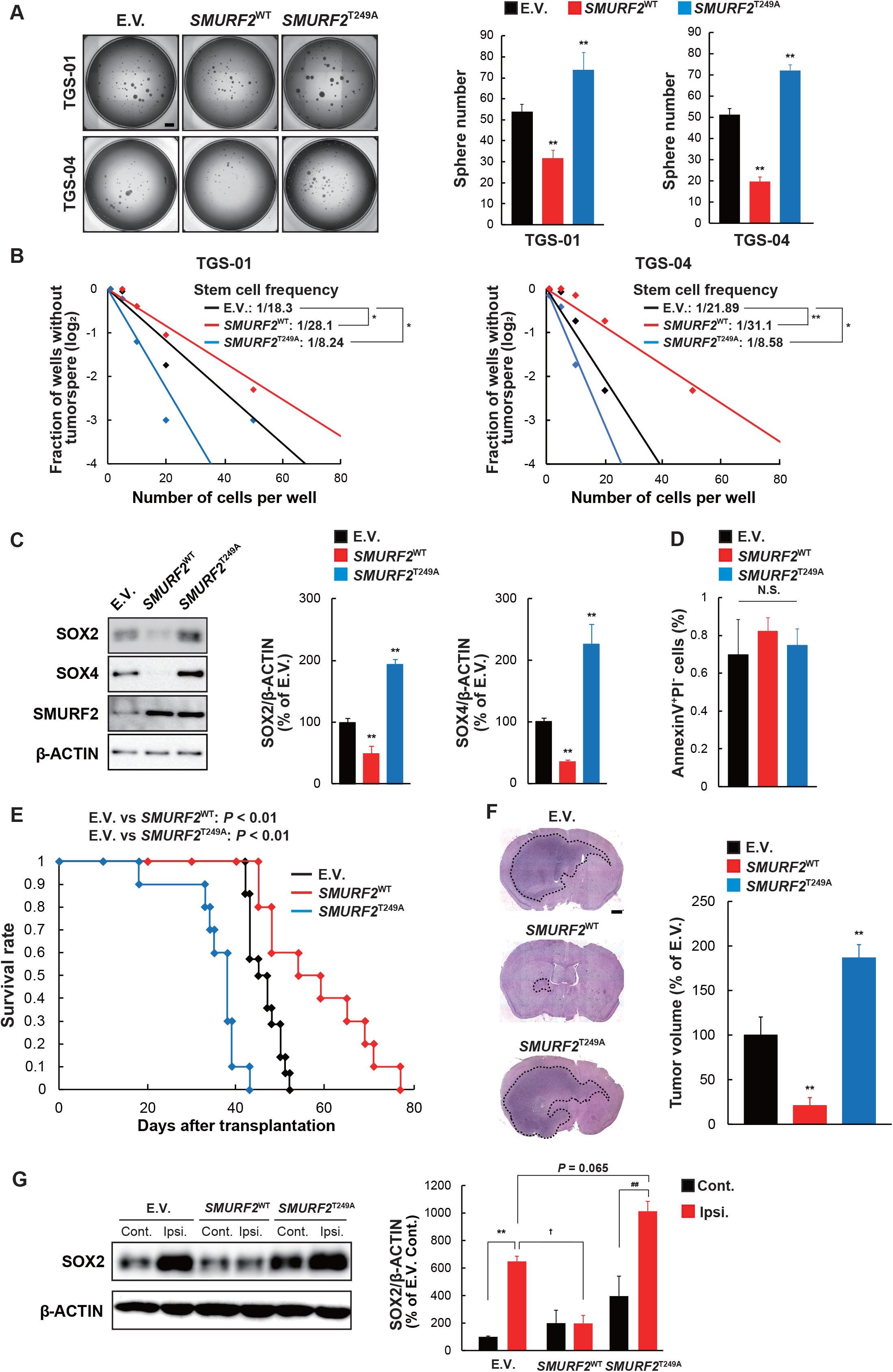
SMURF2^Thr249^ phosphorylation regulates tumor growth and self-renewal of GSCs. TGS-01 and TGS-04 GSCs were infected with *SMURF2*^WT^ or *SMURF2*^T249A^, followed by determination of (A) tumorsphere number (*n*=8), (B) stem cell frequency by *in vitro* limiting dilution assay, (C) protein levels of SOX2, SOX4, and SMURF2 (*n*=3), and (D) cell apoptosis (*n*=3). (E) Development of gliomas after intracranial transplantation of *SMURF2*^WT^- or *SMURF2*^T249A^-infected TGS-01 GSCs. Survival of mice was evaluated by Kaplan-Meier analysis (*n*=14). *P* value was calculated using a log-rank test. (F) Histological analyses of brains dissected at 30 days after intracranial transplantation. Tissue sections were stained with H&E (*n*=5). \**P* < 0.05, \*\**P* < 0.01, significantly different from the value obtained in cells infected with E.V.. (G) Determination of protein levels of SOX2 in the brain of ipsilateral (Ipsi.) side of inoculation and contralateral (Cont.) side at 40 days after intracranial transplantation; β-ACTIN served as a loading control (*n*=3). \*\**P* < 0.01, significantly different from the value obtained in Cont. side inoculated E.V.-infected cells. ^##^*P* < 0.01, significantly different from the value obtained in Cont. side inoculated *SMURF2*^T249A^ -infected cells. ^†^*P* < 0.05, significantly different from the value obtained in Ipsi. side inoculated E.V.-infected cells. N.S., not significant. Values are expressed as the mean ± S.E. and statistical significance was determined using (A, C and F) the one-way ANOVA using the Bonferroni *post hoc* test, (G) Tukey-Kramer test. Scale bar: 1 mm.

We next examined the impact of SMURF2^Thr249^ phosphorylation on the tumorigenic potential of GSCs *in vivo*. Equal numbers of TGS-01 GSCs transduced with either *SMURF2*^T249A^ or *SMURF2*^WT^ were intracranially injected into immunocompromised mice. The inoculation of *SMURF2*^T249A^-infected cells significantly shortened the survival of the mice compared with the inoculation of empty vector (E.V.) -infected cells; conversely, their survival was significantly prolonged by the inoculation of *SMURF2*^WT^-infected cells (Fig. 3E). Moreover, *SMURF2*^T249A^-infected cells generated larger tumors than the control cells, whereas *SMURF2*^WT^-infected cells generated smaller tumors (Fig. 3F). Immunoblotting analysis showed that the SOX2 level was significantly increased in the ipsilateral side than that in the contralateral side after inoculation of E.V.-infected cells and *SMURF2*^T249A^-infected cells (Fig. 3G). The SOX2 level in the ipsilateral side was significantly decreased in mice inoculated with *SMURF2*^WT^-infected cells than that in mice with E.V.-infected cells, whereas it tended to increase in mice inoculated with *SMURF2*^T249A^-infected cells (Fig. 3G). Collectively, these results indicate that SMURF2^Thr249^ phosphorylation could regulate the self-renewal potential and tumorigenicity of GSCs.

### SMURF2^Thr249^ phosphorylation modifies the TGF-β-SMAD2/3 axis by controlling TGFBR stability in GSCs

The self-renewal potential and tumorigenicity of GSCs were activated by inactivating SMURF2^Thr249^ phosphorylation, thus recapitulating GSC phenotypes by *SMURF2* silencing. The phosphorylation of SMURF2^Thr249^ activates its ubiquitin E3 ligase ability to accelerate the proteasomal degradation of SMAD proteins (SMAD1, SMAD2, and SMAD3) in MSCs to control the stemness; furthermore, the TGF-β/SMAD and BMP/SMAD axes play a crucial role in regulating the stemness and tumorigenicity of GSCs through the SMAD pathway (25, 33, 34). We therefore investigated whether SMURF2^Thr249^ phosphorylation could regulate the TGF-β/SMAD and BMP/SMAD axes in GSCs. The protein levels of TGFBR1 and TGFBR2 and the phosphorylation level of SMAD2/3 were significantly increased by *SMURF2*^T249A^; however, these levels were decreased by *SMURF2*^WT^ in TGS-01 GSCs (Fig. 4A). Conversely, the protein levels of BMPR2 and BMPR1A and the phosphorylation level of SMAD1/5/9 were not significantly altered by either *SMURF2*^T249A^ or *SMURF2*^WT^ in TGS-01 GSCs (Fig. 4A). These results indicate that SMURF2^Thr249^ phosphorylation may regulate the TGF-β-SMAD2/3 axis rather than the BMP-SMAD1/5/9 axis in GSCs.

**Figure 4.**
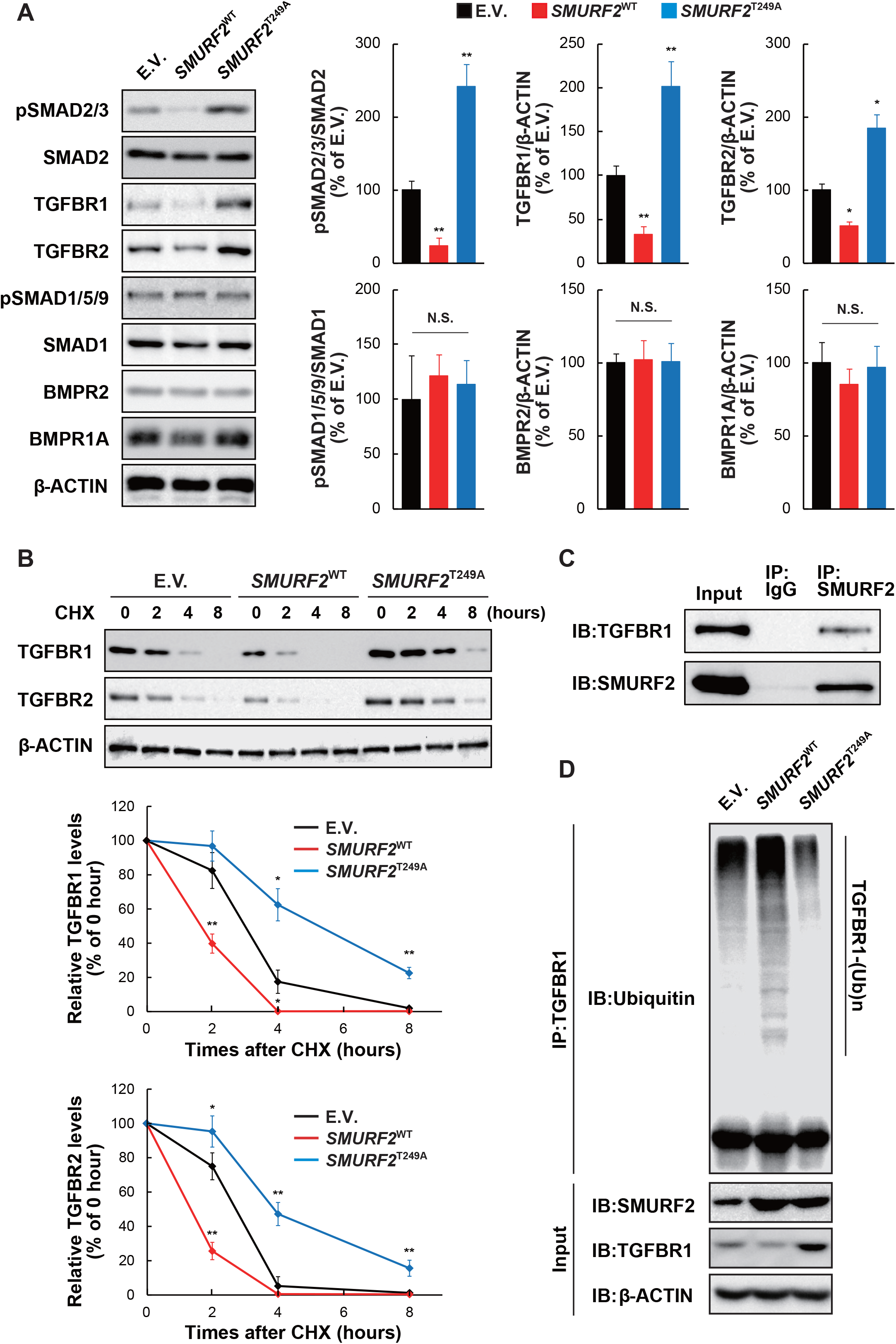
SMURF2^Thr249^ phosphorylation regulates ubiquitin-dependent degradation of TGFBR protein. (A) TGS-01 GSCs were infected with *SMURF2*^WT^ or *SMURF2*^T249A^, followed by determination of protein levels by immunoblotting (*n*=3). (B) TGS-01 GSCs were infected with *SMURF2*^WT^ or *SMURF2*^T249A^, and treated with cycloheximide (CHX) at 50 μg/ml for indicated hours, followed by immunoblotting (*n*=4). (C) Immunoprecipitation assay was performed in TGS-01 GSCs (*n*=3). (D) TGS-01 GSCs were infected with *SMURF2*^WT^ or *SMURF2*^T249A^, and subsequent immunoprecipitation with anti-TGFBR1 antibody, followed by determination of Ubiquitin with anti-Ubiquitin antibody (*n*=3). \**P* < 0.05, \*\**P* < 0.01, significantly different from the value obtained in cells infected with E.V.. N.S., not significant. Values are expressed as the mean ± S.E. and statistical significance was determined using the one-way ANOVA using the Bonferroni *post hoc* test.

To elucidate whether SMURF2^Thr249^ phosphorylation controls TGFBR protein stability through the ubiquitin-proteasome pathway in GSCs, TGS-01 GSCs were treated with cycloheximide (CHX), a protein synthesis inhibitor, and the TGFBR1 and TGFBR2 protein levels were evaluated. The TGFBR1 and TGFBR2 protein levels gradually decreased and became almost undetectable within 8 hours of CHX treatment in E.V.-infected TGS-01 GSCs (Fig. 4B). The enforced expression of *SMURF2*^T249A^ prominently increased the stability of both the TGFBR1 and TGFBR2 proteins whereas the introduction of *SMURF2*^WT^ destabilized their proteins in TGS-01 GSCs (Fig. 4B). We next investigated the role of SMURF2^Thr249^ phosphorylation in SMURF2-dependent TGFBR protein degradation. Firstly, an immunoprecipitation assay revealed that SMURF2 physically interacts with TGFBR1 in TGS-01 GSCs (Fig. 4C). Moreover, endogenous TGFBR1 ubiquitination was markedly elevated by the overexpression of *SMURF2*^WT^, but it was decreased by the enforced infection of *SMURF2*^T249A^ in TGS-01 GSCs (Fig. 4D). These results suggest that SMURF2^Thr249^ phosphorylation decreases the protein stability of TGFBR1 by enhancing its E3 ubiquitin ligase activity, which in turn reduced TGF-β-SMAD2/3 signaling to repress the self-renewal potential and tumorigenicity of GSCs.

### TGFBR1 is a critical target through which SMURF2^Thr249^ phosphorylation can regulate the self-renewal potential and tumorigenicity of GSCs

We next examined whether the activation of TGF-β signaling by TGFBR protein stability could contribute to the regulation of self-renewal potential and tumorigenicity of GSCs by SMURF2^Thr249^ phosphorylation. *TGFBR1* silencing in TGS-01 GSCs significantly attenuated the increased tumorsphere formation and self-renewal potential caused by the introduction of *SMURF2*^T249A^ (Fig. 5A and 5B). Additionally, *TGFBR1* knockdown significantly rescued the shortened duration of survival in mice bearing *SMURF2*^T249A^-infected TGS-01 GSCs, resulting in an increased rate of prolonged survival (Fig. 5C and 5D). Finally, SMURF2^Thr249^ phosphorylation was negatively correlated with the TGFBR1 protein levels in human glioma specimens (Fig. 5E). These results indicate that the phosphorylation of SMURF2^Thr249^ is important for regulating TGFBR1 protein stability to control the self-renewal potential and tumorigenicity of GSCs.

**Figure 5.**
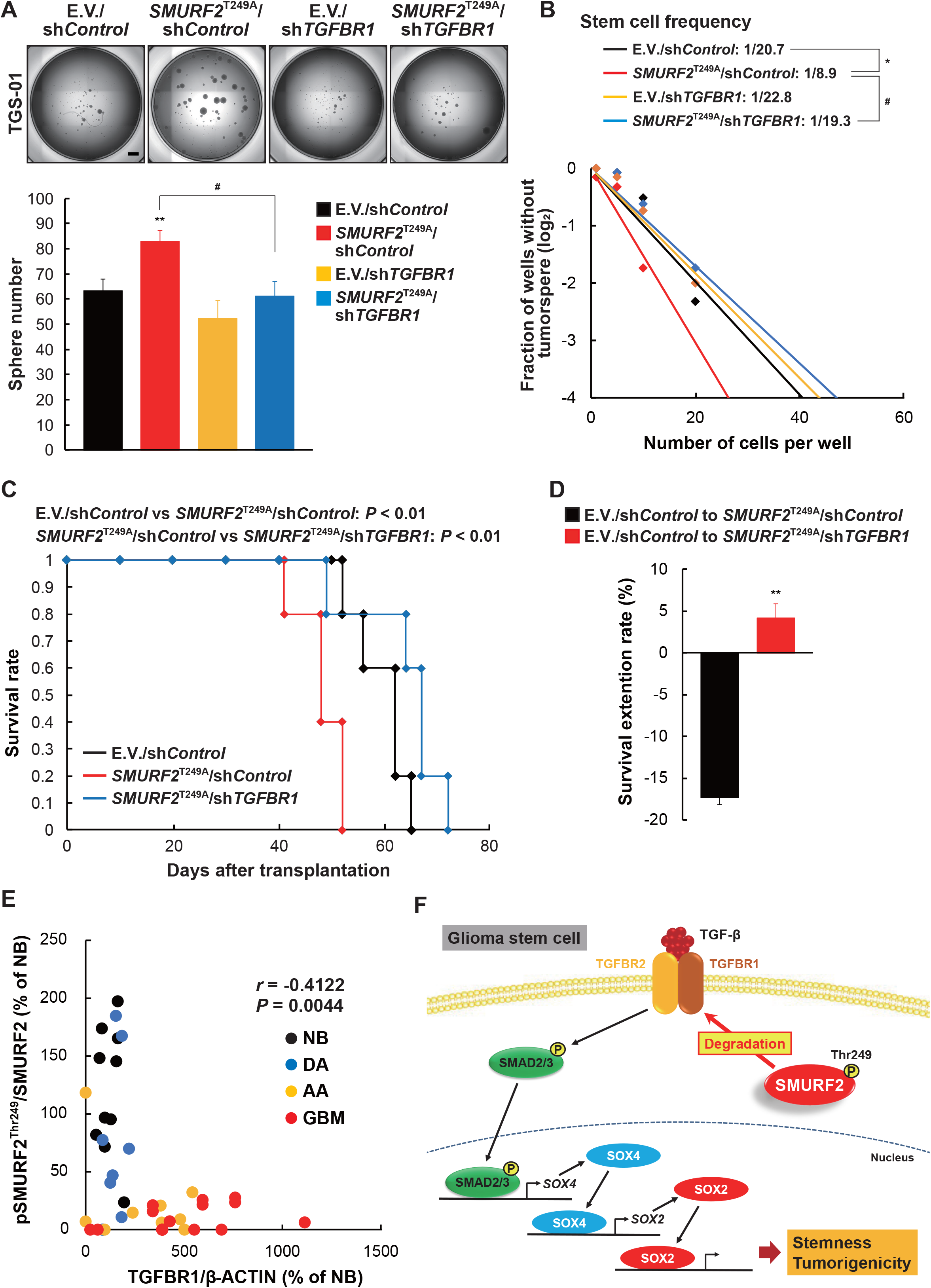
*TGFBR1* silencing restores the promotive effect of *SMURF2*^T249A^ on GSC phenotypes. TGS-01 GSCs were infected with *SMURF2*^T249A^ and/or sh*TGFBR1*, followed by determination of (A) tumorsphere number (*n*=8), (B) stem cell frequency by *in vitro* limiting dilution assay. (C) Development of gliomas after intracranial transplantation of *SMURF2*^T249A^- and/or sh*TGFBR1*-infected TGS-01 GSCs. Survival of mice was evaluated by Kaplan-Meier analysis (*n*=5). *P* value was calculated using a log-rank test. (D) Survival extension rate. (E) Correlation between TGFBR1 and pSMURF2^Thr249^ in glioma samples. (F) Schematic model of the findings of this study. Phosphorylation of SMURF2^Thr249^ enhances ubiquitin-dependent degradation of TGFBR1 protein, which results in the repression of SMAD2/3-SOX4/2 axis, leading to the inhibition of stemness and tumorigenicity of GSCs. (A and B) \**P* < 0.05, \*\**P* < 0.01, significantly different from the value obtained in cells infected with E.V./sh*Control*. ^#^*P* < 0.05, significantly different from the value obtained in cells infected with *SMURF2*^T249A^/sh*Control*. Values are expressed as the mean ± S.E. and statistical significance was determined using the two-way ANOVA using the Bonferroni *post hoc* test. *r*, Pearson’s correlation coefficient. Scale bar: 1 mm.

## Discussion

The E3 ubiquitin ligase activity of SMURF2 is regulated at the posttranslational level, including through phosphorylation (8, 35, 36). SMURF2 activity is inhibited by the phosphorylation at Tyr^314^/Tyr^434^ by c-Src and Ser^384^ by ATM (35, 36). We recently demonstrated that SMURF2^Thr249^ phosphorylation by ERK5 activates its ubiquitin ligase activity and subsequently controls the stemness of MSCs through modulating the SMAD-SOX9 molecular axis, thus contributing to skeletogenesis (10). In this study, SMURF2^Thr249^ phosphorylation controlled the stemness and tumorigenicity of GSCs by modulating the TGFBR-SMAD-SOX4 molecular axis, contributing to gliomagenesis (Fig. 5F), and downregulating SMURF2^Thr249^ phosphorylation in human GBM tissues as well as human GBM patient-derived GSCs. Although further studies should be performed to identify the kinases and phosphatases responsible for controlling SMURF2^Thr249^ phosphorylation in GSCs, our results demonstrated that SMURF2^Thr249^ phosphorylation may be a crucial post-translational modification for modulating the stemness and tumorigenicity of GSCs, thereby suggesting that molecules that modify the activities of kinases and/or phosphatases responsible for SMURF2^Thr249^ phosphorylation could be novel potential GSC-targeting drugs.

SMURF2 is considered to perform a dual role as a promoter and suppressor of tumors by regulating the stability of certain proteins involved in tumorigenesis in cell-dependent and context-dependent manners. SMURF2 interacts with and destabilizes H2AX, which plays a central role in DNA repair and genome stability, in glioma cells (37). *SMURF2* silencing reduces the migration and invasion of breast carcinomas and colorectal cancer (3, 38). Moreover, *SMURF2* is overexpressed in some types of ovarian cancer and breast cancer (4), and high levels of *SMURF2* expression are related to poor prognosis in esophageal carcinomas (39), suggesting that SMURF2 acts as a tumor promoter in certain tumors. Conversely, mouse genetic studies have revealed that *SMURF2* deficiency leads to an increase in the possibility of a wide spectrum of tumors in various tissues and organs including the liver, blood, and lungs in aged mice (40), thus implicating SMURF2 as a potent tumor suppressor. However, the mechanisms underlying SMURF2 activity in human malignancies remain elusive because *SMURF2* is rarely found mutated or deleted in cancers (41). Here, we show that the disruption of *SMURF2* resulted in an enhancement of the self-renewal potential and tumorigenicity of GSCs, which are phenocopied by an inactivation of SMURF2 by a non-phosphorylatable mutant; conversely, the opposite reaction was observed through *SMURF2* overexpression in GSCs. Moreover, SMURF2^Thr249^ phosphorylation was markedly lower in the GBM pathology specimens, accompanied by no marked alteration in the SMURF2 protein level, irrespective of the unknown mechanism of downregulated SMURF2^Thr249^ phosphorylation in GBM patients. Although we should investigate whether SMURF2^Thr249^ phosphorylation has a prognostic value for glioma patients, SMURF2 could exert tumor suppressor functions in glioma pathogenesis, in which SMURF2 activity is controlled by SMURF2^Thr249^ phosphorylation status rather than SMURF2 expression levels.

The functional role of SMURF2 on tumorigenesis has been reported to be connected to its ability to regulate the protein stability of a variety of substrate repertories, in addition to altering the cellular distribution of SMURF2 (3, 40). For example, SMURF2 governs the chromatin organization, dynamics, and genome integrity by controlling the proteasomal degradation or the protein stability of its substrates including RNF20 or DNA topoisomerase IIa (40, 42), which in turn regulate tumorigenesis and tumor progression. Moreover, SMURF2 regulates the stability of pro-oncogenic transcription factors such as KLF5, YY1, and ID1 (43–45), in addition to regulating Wnt/β-catenin oncogenic signaling and KRAS oncoproteins (46–48). Although we show here that SMURF2^Thr249^ phosphorylation plays a crucial role in stemness and tumorigenicity by modulating TGF-β signaling through the ubiquitin-proteasome-dependent degradation of TGFBR proteins in GSCs, it should be emphasized that additional molecular mechanisms might be involved in the control of tumorigenicity in GSCs by SMURF2^Thr249^ phosphorylation.

In conclusion, SMURF2^Thr249^ phosphorylation plays a crucial role in glioma pathogenesis by modulating TGF-β/SMAD signaling in GSCs. To our knowledge, this is the first preclinical study to investigate the functional role of SMURF2 on the function of cancer stem cells *in vivo*. Our findings improve our understanding of the molecular mechanism underlying the maintenance of the stemness and tumorigenicity of GSCs and suggest that SMURF2^Thr249^ phosphorylation status could represent a novel target for drug development to treat not only gliomas but also malignant tumors associated with the aberrant expression or function of TGF-β signaling in humans.

## Materials and Methods

### Cell culture and reagents

HEK293T cells were purchased from RIKEN BRC (#RCB2202). HEK293T cells were cultured in Dulbecco’s modified Eagle’s medium (DMEM) (FUJIFILM Wako Pure Chemical #043-30085) supplemented with 10% fetal bovine serum. Human patient-derived GBM cell lines TGS-01 and TGS-04 were established as described previously (25). The use of these human materials and protocols were approved by the Ethics Committees of Gifu Pharmaceutical University, Kanazawa University, and the University of Tokyo. These cells were cultured in neurosphere medium containing DMEM/F12 (FUJIFILM Wako Pure Chemical #048-29785) supplemented with GlutaMAX (Gibco #35050061), B27 supplement minus vitamin A (Gibco #12587010), 20 ng/ml recombinant human epidermal growth factor (FUJIFILM Wako Pure Chemical #059-07873) and 20 ng/ml recombinant human basic fibroblast growth factor (FUJIFILM Wako Pure Chemical #064-04541). These cells were differentiated in adherent culture medium containing DMEM supplemented with 10% fetal bovine serum for 7 days.

### Surgical specimens

A total of 46 primary glioma tissues were obtained from patients who underwent surgical removal of tumor. The specimens were reviewed and classified according to WHO criteria (49). Nonneoplastic healthy brain tissues adjacent to tumors were acquired. The tissues were homogenized in lysis buffer. All experiments were approved by the local Institutional Review Board of Kanazawa University (No. 2509) and all study participants provided written informed consent.

### Immunoblotting analysis

Cells were solubilized in lysis buffer (10 mM Tris-HCl, 150 mM NaCl, 0.5 mM EDTA, 10 mM NaF, 1% Nonidet P-40, pH 7.4) containing protease inhibitor cocktail. Samples were then subjected to SDS-PAGE, followed by transfer to polyvinylidene difluoride (PVDF) membranes and subsequent immunoblotting. The primary antibodies used were, anti-p-Smurf2^Thr249^ (#J1683BA260-5, 1:2000) (GenScript), anti-Phospho-Smad2 (Ser465/467) (#3101, 1:1000), anti-Smad2 (#5339, 1:1000), anti-TGF-β Receptor I (#3712, 1:1000), anti-TGF-β Receptor II (#79424, 1:1000), anti-Sox2 (#3579, 1:1000), anti-p-Smad1 (Ser463/465)/5 (Ser463/465)/9 (Ser465/467) (#13820, 1:1000), anti-Smad1 (#9512, 1:1000), anti-BMPR2 (#6979, 1:1000) and anti-Ubiquitin (#3936, 1:1000) (Cell Signaling Technologies), anti-β-actin (#sc-47778, 1:2000) (Santa Cruz Biotechnology), anti-Sox4 (#AB5803, 1:1000) (EMD Millipore), anti-BMPR1A (#ab174815, 1:1000) and anti-SMURF2 (#ab94483, 1:1000) (Abcam). The primary antibodies were diluted with blocking solution (5% skim milk). The custom polyclonal p-Smurf2^Thr249^ antibody was generated (#J1683BA260-5) (GenScript). Briefly, two rabbits were injected with KLH-conjugated p-Smurf2^Thr249^ epitope, representing amino acids 244-258, emulsified in Freund’s complete adjuvant, and then boosted 3 times at 14-day intervals with p-Smurf2^Thr249^ epitope. The images were acquired using ChemiDoc Touch Imaging System (Bio-Rad). Quantification was performed by densitometry using ImageJ.

### Tumorsphere formation assay and *in vitro* limiting dilution assay

For sphere formation assay, single cell suspensions were prepared using StemPro Accutase (Gibco, #A1110501) and filtered through a 70 μm cell strainer (BD Biosciences). Cells were then plated in 96-well Costar ultra-low attachment plate (Corning) at 2 × 10^3^ cells per well with neurosphere medium mixed with 1% methylcellulose. Tumorsphere number were measured on day 7. For *in vitro* limiting dilution assay, cells were plated in 96-well plate at 1, 5, 10, 20, 50, 100 or 200 cells per well, with 10 replicates for each cell number. The presence of tumorspheres in each well was examined on day 7. Cell images were captured using a BZ-X810 fluorescence microscope (Keyence) and analyzed for quantitating sphere numbers and sizes using BZ-X810 Analyzer software (Keyence). Limiting dilution assay analysis was performed using online software (http://bioinf.wehi.edu.au/software/elda/). Sphere formation was estimated by scoring the number of spheres larger than 50 μm.

### Orthotopic xenograft model of GSC-derived GBM and histology

Orthotopic xenograft model of GSC-derived GBM was generated by transplantation of 5 × 10^4^ TGS-01 GSCs into the brain of 4-week-old female nude mice (BALB/cSlc-nu/nu, SLC, Shizuoka, Japan). Briefly, a small burr hole was drilled in the skull 0.5 mm anterior and 2.0 mm lateral from bregma with a micro drill, and dissociated cells were transplanted at a depth of 3 mm below the dura mater. Mice were sacrificed at the indicated time points or upon occurrence of neurological symptoms. Mouse brains were fixed with 4% paraformaldehyde solution, embedded in paraffin, and then sectioned at a thickness of 5 μm. Sections were stained with Hematoxylin and Eosin (H&E). The sections were captured using a BZ-X810 fluorescence microscope (Keyence). All animal experiments were approved by the Committees on Animal Experimentation of Gifu Pharmaceutical University and Kanazawa University and performed in accordance with the guidelines for the care and use of laboratory animals. The numbers of animals used per experiment are stated in the figure legends.

### Generation of lentiviral vectors and infection

The lentiviral *SMURF2* mutant vector was previously generated (10). The oligonucleotides for *SMURF2* short hairpin RNA (shRNA) were synthesized (Supplementary Table), annealed, and inserted into the mCherry vector, and the shRNA vector for *TGFBR1* was obtained from Sigma (SHCLNG-NM_004612, TRCN0000196326). These vectors were then transfected into HEK293T cells using the calcium carbonate method. Virus supernatants were collected 48 h after transfection and cells were then infected with viral supernatant for 24 h.

### Flow cytometry

Cells were dissociated into single cells with StemPro Accutase (Gibco). Apoptosis assay was conducted using FITC-Annexin V Apoptosis Detection kit (BD Biosciences, #556547) and analyzed by BD FACS Verse and BD FACSuite software.

### Immunoprecipitation (IP) assay

Cells were solubilized in lysis buffer (10 mM Tris-HCl, 150 mM NaCl, 0.5 mM EDTA, 10 mM NaF, 1% Nonidet P-40, pH 7.4) containing protease inhibitor cocktail. Samples were incubated with an antibody in lysis buffer for 24 h at 4 °C and subsequent IP with protein G-Sepharose. Immunoprecipitates were washed three times with lysis buffer and boiled in SDS sample buffer. Samples were then separated by SDS-PAGE, followed by transfer to PVDF membranes and subsequent immunoblotting.

### Bioinformatics

Gene expression data from the Cancer Genome Atlas (TCGA) project was obtained and analyzed using GlioVis database (http://gliovis.bioinfo.cnio.es/).

### Statistical analysis

Unless otherwise specified, Student’s *t*-test and one-way ANOVA *post hoc* Bonferroni test were used for statistical significance. Throughout this study, *P* < 0.05 were considered statistically significant. For correlation analysis, we calculated Pearson’s correlation coefficient.

## Acknowledgements

This work was supported in part by the Japan Society for the Promotion of Science (20H03407 to E.H.); and a grant from Japan Research Foundation for Clinical Pharmacology (to E.H.).

